# The mycobacterial glycoside hydrolase LamH enables capsular arabinomannan release and stimulates growth

**DOI:** 10.1101/2023.10.26.563968

**Authors:** Aaron Franklin, Abigail J. Layton, Todd Mize, Vivian C. Salgueiro, Rudi Sullivan, Samuel T. Benedict, Sudagar S. Gurcha, Itxaso Anso, Gurdyal S. Besra, Manuel Banzhaf, Andrew L. Lovering, Spencer J. Williams, Marcelo E. Guerin, Nichollas E. Scott, Rafael Prados-Rosales, Elisabeth C. Lowe, Patrick J. Moynihan

## Abstract

Mycobacterial glycolipids are important cell envelope structures that drive host-pathogen interactions. Arguably, the most important amongst these are lipoarabinomannan (LAM) and its precursor, lipomannan (LM), which are both trafficked out of the bacterium to the host via unknown mechanisms. An important class of exported LM/LAM is the capsular derivative of these molecules which is devoid of its lipid anchor. Here, we describe the identification of a glycoside hydrolase family 76 enzyme that we term LamH which specifically cleaves α-1,6-mannoside linkages within LM and LAM, driving its export to the capsule releasing its phosphatidyl-*myo*-inositol mannoside lipid anchor. Unexpectedly, we found that the catalytic activity of this enzyme is important for efficient exit from stationary phase cultures where arabinomannan acts as a signal for growth phase transition. Finally, we demonstrate that LamH is important for *Mycobacterium tuberculosis* survival in macrophages. These data provide a new framework for understanding the biological role of LAM in mycobacteria.

## Introduction

The bacterial cell envelope includes a diverse array of molecules that in addition to contributing to basic biology, act as messengers that help shape or report on their environmental niche. In the context of host-pathogen interactions, these molecules are typically chemically distinct, marking the bacteria as different from their host. For mycobacterial pathogens, their unusual cell envelope includes a wide range of such structures that act in concert to manipulate the host immune response. The core elements of this are the mycolyl-arabinogalactan-peptidoglycan complex, and lipoglycans such as lipomannan (LM) and lipoarabinomannan (LAM) (Fig. 1a)^1^.

**Figure 1.**
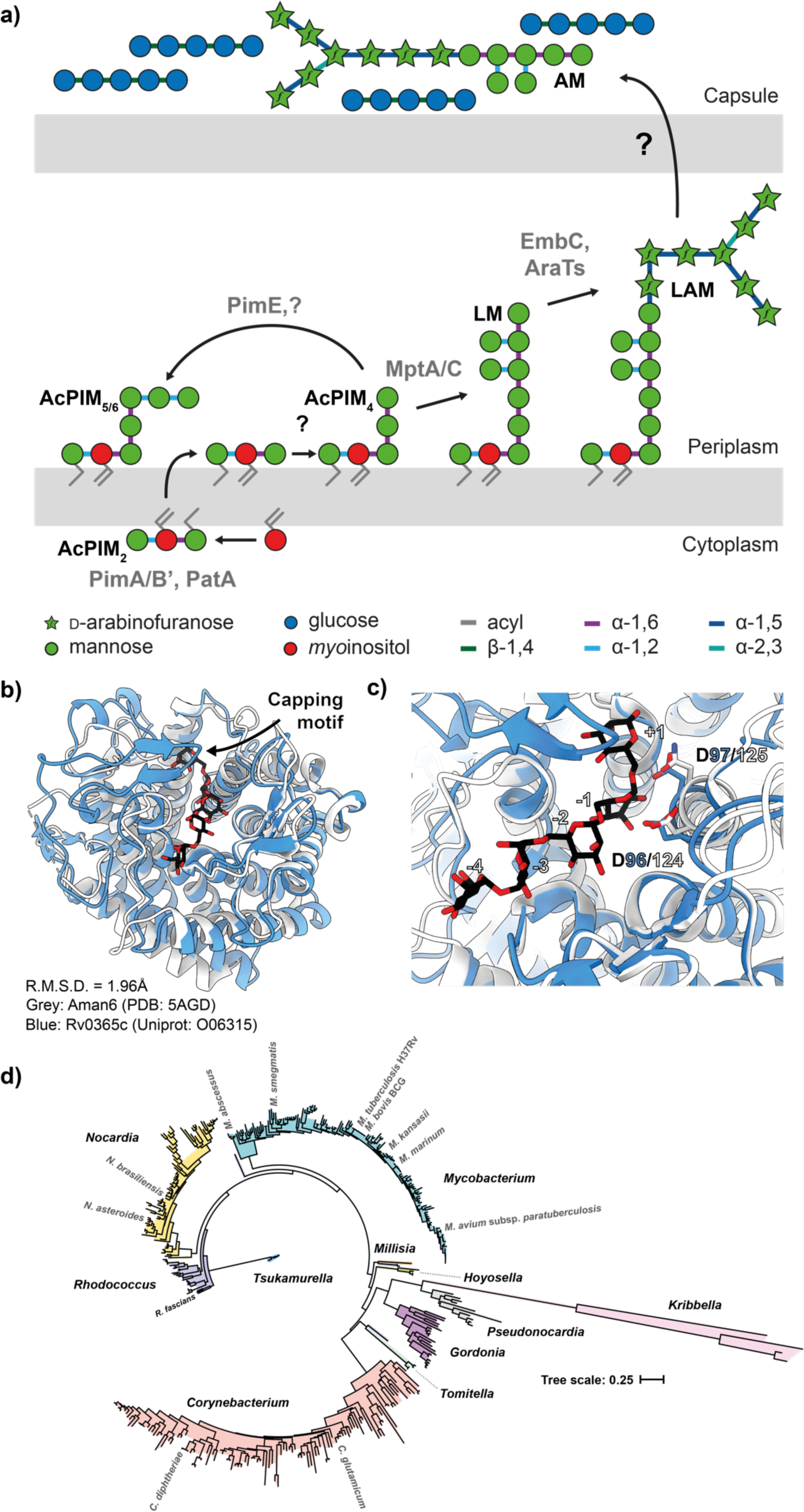
Biosynthesis of mannosylated glycolipids in mycobacteria. **a)** The biosynthetic pathway for LM and LAM results in three primary products, PIMs, LM and LAM. The degree of acylation for each product can vary, as can secondary modifications on LM and LAM. An unknown process drives expression of AM and M on the surface of the bacteria as part of their capsule. A simplified structure of LM/LAM is presented for clarity. Grey text indicates glycolipid species. **b)** Ribbon diagram of Aman6 (PDB:5AGD; grey) in complex with ɑ-1,6 mannotetraose aligned in ccp4i Superpose with the AlphaFold 2.0 predicted structure for LamH (Rv0365c) associated with its Uniprot entry (O06315; blue). **c)** The confirmed catalytic residues in Aman6 (D124/D125; D125N mutant in 5AGD; grey) are conserved in LamH (D96/D97; blue). **d)** Phylogenetic tree of GH76 enzymes from the Mycobacteriales. Genomes for all available members of the Mycobacteriales were used to generate a custom BLAST database in Geneious Prime 2023.1.1. LamH was used as a query in a BLAST search of this database, yielding high-confidence homologs from all species. This list of proteins was submitted to NGPhylogeny.fr using the PHYML/OneClick tool, the tree was then manually coloured to identify individual genera.

LAM and LM are important to the pathogenesis of mycobacteria and form the basis of many diagnostic platforms^2^. They are required to control acidification of the phagosome and are a ligand for the C-type lectins Dectin-2 and DC-Sign^3,4^. Through decades of careful analysis, the structure of LM and LAM has been revealed to include three structural domains. The first is a phosphatidyl-*myo*-inositol anchor, believed to be biosynthetically derived from phosphatidyl-*myo*-inositol mannosides (PIMs)^5^. Attached to this is a large mannan moiety, with an α-1,6 mannose backbone of approximately 13 residues branched with α-1,2 mannose decorations^6^. A large, branched D-arabinofuranose domain is connected to this mannan backbone, which is far more complex and includes multiple non-stoichiometric acyl modifications^7–9^. Much of the immunogenicity of LAM has been associated with the final, capping structures on the molecule, most notably in the form of α-1,2-linked mannose caps and an unusual methylthioxylose residue in *M. tuberculosis*^10–13^. While broadly similar across species, the precise structure of LM and LAM associated with the cytoplasmic membrane can vary between species and strains with substantial impacts on the host response^14^. Furthermore, while the chemical analysis of LAM derived from bacteria has proved incredibly useful, the structures of the macromolecules that are ultimately secreted and perceived by the host are not well defined. LM and LAM have also been proposed to play important roles in the basic biology of the bacteria, contributing to proper septal formation in a manner analogous to lipoteichoic acids in some Gram-positive bacteria^15,16^. Nonetheless, much of the biology of LM and LAM remains to be uncovered.

Amongst the pathways that require further study is the modification and release of LM and LAM. Release of these molecules may happen passively, as is the case during cell wall degradation because of division and wall remodelling, or actively as the bacteria secrete effectors into the host^17,18^. In the context of mycobacteria, we recently described the identification of a new family of glycoside hydrolases (GHs) responsible for the release of D-arabinan fragments into the surrounding environment^19^. The fate of these fragments is largely unknown, however at least some of these molecules end up in the mycobacterial capsule^20^. The composition of this capsule varies substantially amongst mycobacterial species, but in *M. tuberculosis* the carbohydrate component has been reported to be comprised of 80% α-glucan and approximately 20% arabinomannan (AM) and mannan^20^. The latter two of these are believed to be derived from LM and LAM, however no mechanism for this has been established.

To address this gap in our understanding, we have identified LamH (Rv0365c) as the enzyme responsible for release of the carbohydrate domain of LM and LAM to the capsule of mycobacteria. We show that the enzyme is highly specific for the attachment point between the lipid anchor and the carbohydrate domain of LM and LAM and plays an important role in the biology of the bacilli. Loss of LamH results in a prolonged lag-phase, which is coupled with an accumulation of LM and LAM and a down-regulation in the production of each of these molecules. Our data also shows that during the lag phase, AM can act as a signalling molecule, whose accumulation helps the bacteria trigger transition to exponential growth. Finally, we show that this protein enables the correct processing and display of capsular AM and mannan, and that knock-down of this protein in *M. tuberculosis* results in a decrease in pathogenesis. Altogether our data provide a new perspective on LM and LAM and their release into the extracellular environment.

## Results

### Mycobacteria encode a single predicted family GH76 enzyme

Prior work has identified the GH76 family as a large group of enzymes that can either degrade, or in a subset of these enzymes, generate through transglycosylation, α-1,6 mannoside linkages^21–24^. For example, we have previously shown that GH76 enzymes from the Bacteroidota are essential for the cleavage of α-1,6 mannoside linkages in fungal mannan in the human gut^23^. We used the experimental structure of Aman6 (PDB:5AGD), a known GH76 family member, as a search model in Foldseek and limited the taxon to *M. tuberculosis* H37Rv^25^. This search yielded a single high confidence result, supporting identification Rv0365c as the sole predicted family GH76 enzyme encoded within the *M. tuberculosis* genome. While broadly similar (LSQKab calculated R.M.S.D. = 3.49A), the mycobacterial enzyme is predicted to possess an additional β-hairpin cap covering the active site (Fig. 1b,c)^26^. We then sought to determine if homologs of Rv0365c are conserved amongst mycobacteria. We identified Rv0365c homologs using a custom BLAST database comprised of representative genomes for members of the Mycobacteriales and used NGPhylogeny.fr to reconstruct a phylogenetic tree (Fig. 1d)^27,28^. Rv0365c homologs were conserved in all representative species, suggesting they participate in an evolutionarily conserved process.

### Rv0365c specifically cleaves α-1,6 mannoside linkages

We next wanted to determine if Rv0365c possessed GH76-like catalytic activity. LM and LAM are complex substrates, making them unsuited to determining the precise linkage specificity of the enzyme. To circumvent this, we purified mannan from three *Saccharomyces cerevisiae* strains lacking glycosyltransferases that produce three distinct mannans of decreasing complexity (Extended Data Fig. 1a)^23^. Each of these have a backbone of α-1,6 linked mannose, while Mnn5 and Mnn1 have additional α-1,2-linked mannose decorations and extensions respectively. Our initial assays with purified Rv0365c on these substrates yielded no observable reaction products (Fig. 2a). Many endo-acting GHs have been shown to preferentially cleave shorter substrates, and so we pre-digested the yeast mannans with BT3792 which produces shorter oligosaccharides. Incubation of pre-processed mannans with Rv0365c revealed an endo-like reaction profile exclusively from the material derived from the strain Mnn2 (Fig. 2b). These data demonstrate α-1,6 mannanase activity for Rv0365c and indicate that the enzyme is unable to cleave substrates with α-1,2-linked mannose decorations. To further confirm this finding, we isolated an ɑ-1,6-mannotetraose oligosaccharide from this mixture, and digested it with Rv0365c, supporting its designation as an endo-ɑ-1,6-mannanase (Fig. 2c). Taken together we conclude that Rv0365c is a member of the GH76 family and is specific for undecorated α-1,6 mannan.

**Figure 2.**
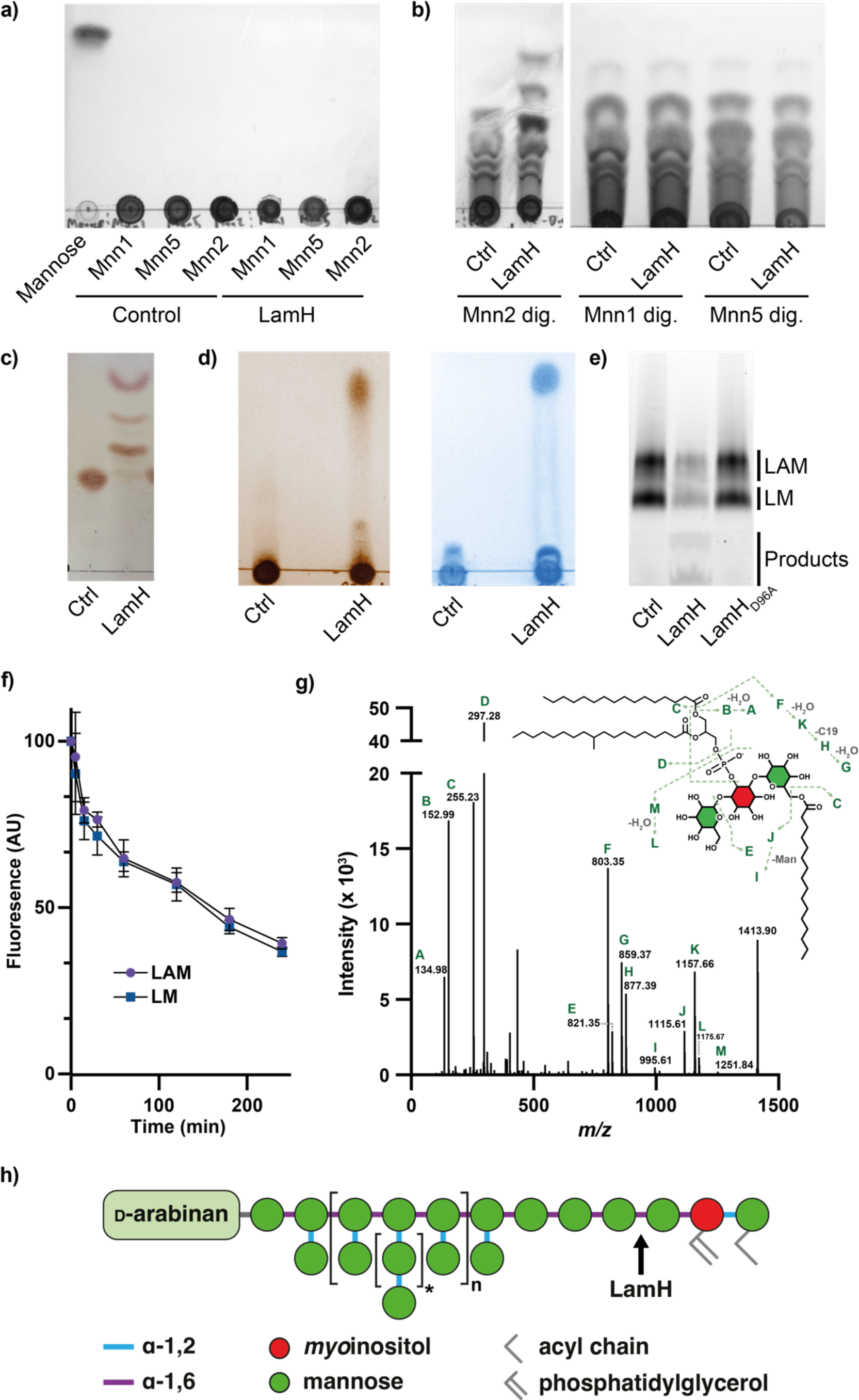
LamH (Rv0365c) is a GH76 family enzyme that cleaves LM and LAM to produce AcPIM_2_. **a)** Yeast mannan substrates were incubated with LamH for 16 h at 37 °C, separated by TLC, and stained with orcinol. **b)** Yeast mannans were incubated with BT3792 for 3 h at 37 °C and the heat inactivated at 100 ℃ for 10 min. The digested substrates were then incubated with LamH for 16 h at 37 ℃, separated by TLC, and stained with orcinol. **c)** LamH was incubated with purified ɑ-1,6 mannotetraose and the reaction products were analysed by TLC, demonstrating endo-activity. **d)** LamH was incubated with mixed LM/LAM and the reaction product was analysed by TLC. Duplicate TLCs were stained with either orcinol or phosphomolybdic acid which indicate the presence of carbohydrate and lipid respectively revealing a small product migrating towards the top of the TLC. **e)** Reaction products were analysed by SDS-PAGE with Pro-Q Emerald staining. A parallel reaction was conducted with the predicted catalytic-null construct LamH_D96A_. **f)** LamH activity against LM and LAM was compared by separating reaction products at the indicated time points and quantifying the LM and LAM fluorescence in ProQ-Emerald stained SDS-PAGE gels. **g)** The small glycolipid identified in Fig. 2b was isolated and analysed by MS/MS. The fragmentation pattern is consistent with an AcPIM_2_ species. **h)** Schematic diagram of LM/LAM based on Angala *et al.* 2020 with the site of LamH activity indicated.

### Rv0365c cleaves the lipid anchor from LM and LAM

In mycobacteria, α-1,6-linked mannan has only been described in regions of LM, LAM, and their presumed capsular derivatives. We therefore hypothesised that Rv0365c would specifically degrade LM and LAM. To test this, we purified these lipoglycans from *Mycobacterium bovis* BCG Danish and incubated them with the enzyme. The reaction products were analysed by TLC, using a solvent system that would retain the carbohydrate domain at the origin, and separate any released glycolipids. As shown in Fig. 2d, Rv0365c released a low molecular weight product from LM and LAM, which stained positive for both carbohydrates and lipid. We also separated the LM/LAM reaction products by SDS-PAGE followed by ProQ-Emerald staining, demonstrating activity on both LM and LAM (Fig. 2e). Incubation of the enzyme with isolated PIMs showed no detectable product formation, suggesting the enzyme cannot process PIM_5_ or PIM_6_ substrates, which is consistent with its inability to degrade α-1,2-linked mannose decorated substrates (Extended Data Fig. 1b).

Supported by comparison to Aman6 (Fig. 1c) we predicted that D96 and D97 (numbering relative to *M. tuberculosis* H37Rv genomic annotation) are the catalytic acid/base and nucleophile residues in this enzyme. We therefore generated the D96A mutant, which was catalytically inactive (Fig. 2e). We next sought to determine if the enzyme preferentially degraded LM or LAM. We quantified the degradation of both species over time by incubating the enzyme with LM/LAM and then analysing a time-course of the reaction products by SDS-PAGE (Extended Data Fig. 1c). This revealed that LM and LAM bands decrease at the same rate (Fig. 2f) and that the enzyme exhibits no preference for either substrate. Prior work indicates that there are on average between 2 (*M. tuberculosis*) and 7 (*Mycobacterium smegmatis*) α-1,6-linked mannose residues immediately attached to the *myo*-inositol anchor which lack α-1,2 decorations^7,29^. Given the many possible cleavage sites available for Rv0365c, we next sought to identify the limit glycolipid product formed by exhaustive Rv0365c digestion of LAM. The sole glycolipid product was analysed by MS/MS (Fig. 2g), revealing the product to be AcPIM_2_. The site of cleavage on LAM is illustrated in Fig. 2h, using the most recent structural proposal of LAM from *M. tuberculosis* as a template^6^. From these data we conclude that Rv0365c degrades LM and LAM, releasing the carbohydrate domain from the AcPIM_2_ lipid anchor and have renamed the protein LamH (<u>L</u>ipo<u>a</u>rabino<u>m</u>annan <u>H</u>ydrolase).

### LamH drives production of capsular (arabino)mannan

Our biochemical data support the hypothesis that LamH is responsible for generating the capsular products of LM and LAM. To test this, we utilised a transposon mutant within *lamH* (BCGDAN_0378) in *M. bovis* BCG Danish, hereafter referred to as Δ*lamH*^30^. This species is 99% genetically identical to *M. tuberculosis*, having lost elements predominantly related to pathogenesis and is generally an excellent model system for *M. tuberculosis* envelope biogenesis and turnover. We first sought to determine if the loss of *lamH* had any impact on the levels of LM and LAM in the mutant. We observed accumulation of approximately 30% more LM/LAM in the Δ*lamH* strain compared to the wild-type strain at mid-exponential growth (Fig. 3a,b, Extended Data Fig. 2a). Consistent with the biochemical data showing no preference for cleavage of LM and LAM by LamH, the change was approximately equal in magnitude for both LM and LAM. We generated a complementation plasmid in the pMV306 plasmid including the *lamH* open reading frame and 300 upstream bases, to include the likely promoter elements. Introduction of this *lamH* locus at the L5 site fully complemented the over-production of LM/LAM in the complemented mutant, but a catalytic null variant of this construct phenocopied the Δ*lamH* strain (Fig. 3a). We also deleted *lamH* (MSMEG_0740) in *M. smegmatis* mc^2^155 and analysed the LM/LAM composition in this strain, which gave results consistent with that of *M. bovis* BCG Danish (Extended Data Fig. 3 a,b). Collectively, these data support the hypothesis that LamH is responsible for maintaining LM/LAM levels within the cell.

**Figure 3.**
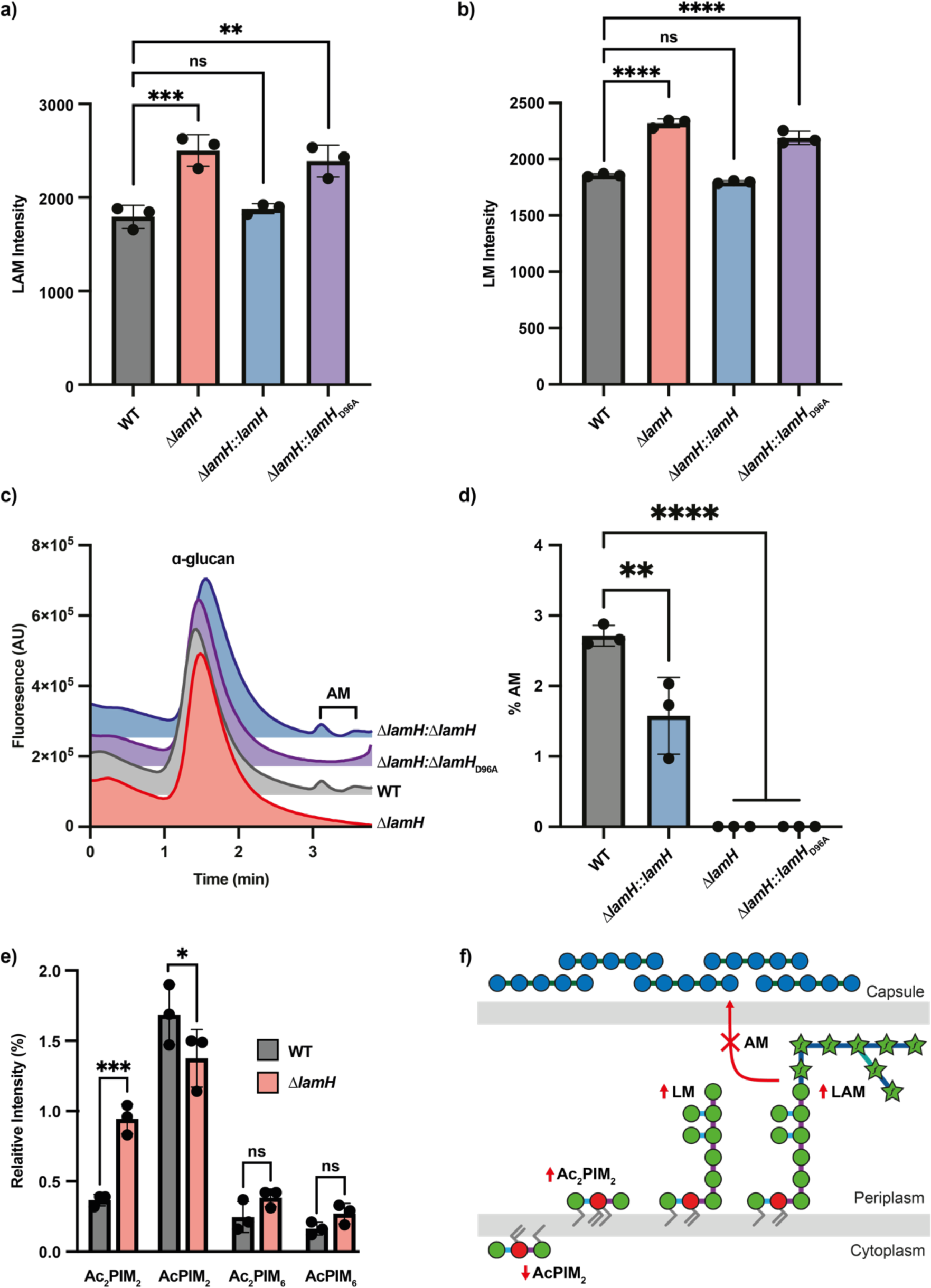
A Δ*lamH* strain retains LAM in the cytoplasmic membrane and does not produce capsular AM. **a)** LM/LAM was harvested from the indicated *M. bovis* BCG strains at OD_600_ = 0.6 and analysed by SDS-PAGE followed by ProQ Emerald staining (Extended Data Fig. 1). Total fluorescence of combined LAM and LM for three biological replicates is presented with error bars indicating standard deviation. **b)** The indicated strains were grown on 7H10 agar and capsular material was isolated and normalised based on wet mass. Glycans were end-labelled with 2-AB and separated by size exclusion chromatography with fluorescence detection (Ex= 320 nm Em = 420 nm). Peaks were identified by digestion with appropriate enzymes. **c)** The α-glucan and AM peaks from Fig. 3c were integrated using Chromeleon 7.2 and the relative amount of AM was computed by dividing the total AM peak area by the sum of the α-glucan and AM peak areas. **d)** The indicated *M. bovis* BCG strains were grown in 7H9 in the presence of (1-^14^C) acetic acid, sodium salt until OD_600_ = 0.6. Polar lipid extracts were analysed by two-dimensional TLC and visualised on exposure X-ray film by autoradiography. TLCs were annotated as per Driessen *et al.,* (2009). Relative intensity of each PIM species is reported. **f)** Schematic summary of phenotypic changes due to loss of *lamH.* Significance was determined with a one-way ANOVA with Tuckey’s post-hoc test. *P < 0.05, **P < 0.01, ***P < 0.001, ****P < 0.0001.

To investigate if LamH is responsible for the production of capsular (arabino)mannan, we examined levels of AM in the capsule of wild-type and mutant strains. Capsular polysaccharides from these strains were isolated and labelled with 2-aminobenzamide, then separated by size-exclusion chromatography, which allows the separation of α-glucan (∼100 kDa) from AM (∼14 kDa)^20^. As shown in Fig. 3c, while AM was detected in wildtype, no AM was detected in the capsule of the Δ*lamH* strain, and capsular AM production was rescued in the complemented strain. The production of capsular AM required the enzyme be catalytically active (Fig. 3c). As 2-AB labelling instals a single label at each reducing end of the glycans, this allows calculation of the ratio of reducing ends of α-glucan to AM which can be used as a proxy for capsular composition. This analysis indicated that the complemented mutant produced significantly less AM than the wild type (Fig. 3d), showing that complementation is imperfect, and suggesting that expression of the *lamH* gene is altered when located distally. We also analysed the PIM composition of the wild-type and mutant bacteria (Fig. 3e, Extended Data Fig. 2b). These data indicate that in response to *lamH* deletion, the bacteria produce less AcPIM_2_ but significantly more Ac_2_PIM_2_. It was recently demonstrated that acylation of PIMs can occur as a response to membrane stress, which suggests that increased abundance of LM/LAM triggers a membrane stress response^31^. Taken together these data are consistent with the hypothesis that LamH drives production of capsular AM by cleaving LAM. *Mycobacteria respond to the lack of LamH activity by down-regulating LAM biogenesis* Given that Δ*lamH* bacteria were unable to degrade existing LM/LAM, we were surprised that they did not accumulate even more of these materials. To better understand why this might be the case we conducted whole-cell proteomics of mid-exponential bacteria (Supplementary Table 1). The Δ*lamH* strain showed a pronounced reduction in abundance of several enzymes in the LM/LAM biogenesis pathway, suggesting that the bacteria respond to the accumulation of LM/LAM by reducing synthesis (Fig. 4a,b). While not all biosynthetic enzymes were observed, even within wild type, levels of arabinosyltransferases associated with LAM biogenesis (AftB, AftC, AftD) were observed to be significantly reduced or undetectable within Δ*lamH* (Supplementary Figure 4)^32–35^. Similarly, the levels of EmbC and MptA, which are arabinosyl- and mannosyl-transferases associated with LAM biogenesis were both reduced, however not significantly so^36,37^. In contrast, the expression of PimA and PatA, which are involved in both PIM and LM/LAM biogenesis are unchanged (Extended Data Fig. 4)^38–41^. Taken together, this provides evidence that the levels of LM/LAM are tightly regulated by the bacteria and that LamH is involved in maintaining LM/LAM homeostasis.

**Figure 4.**
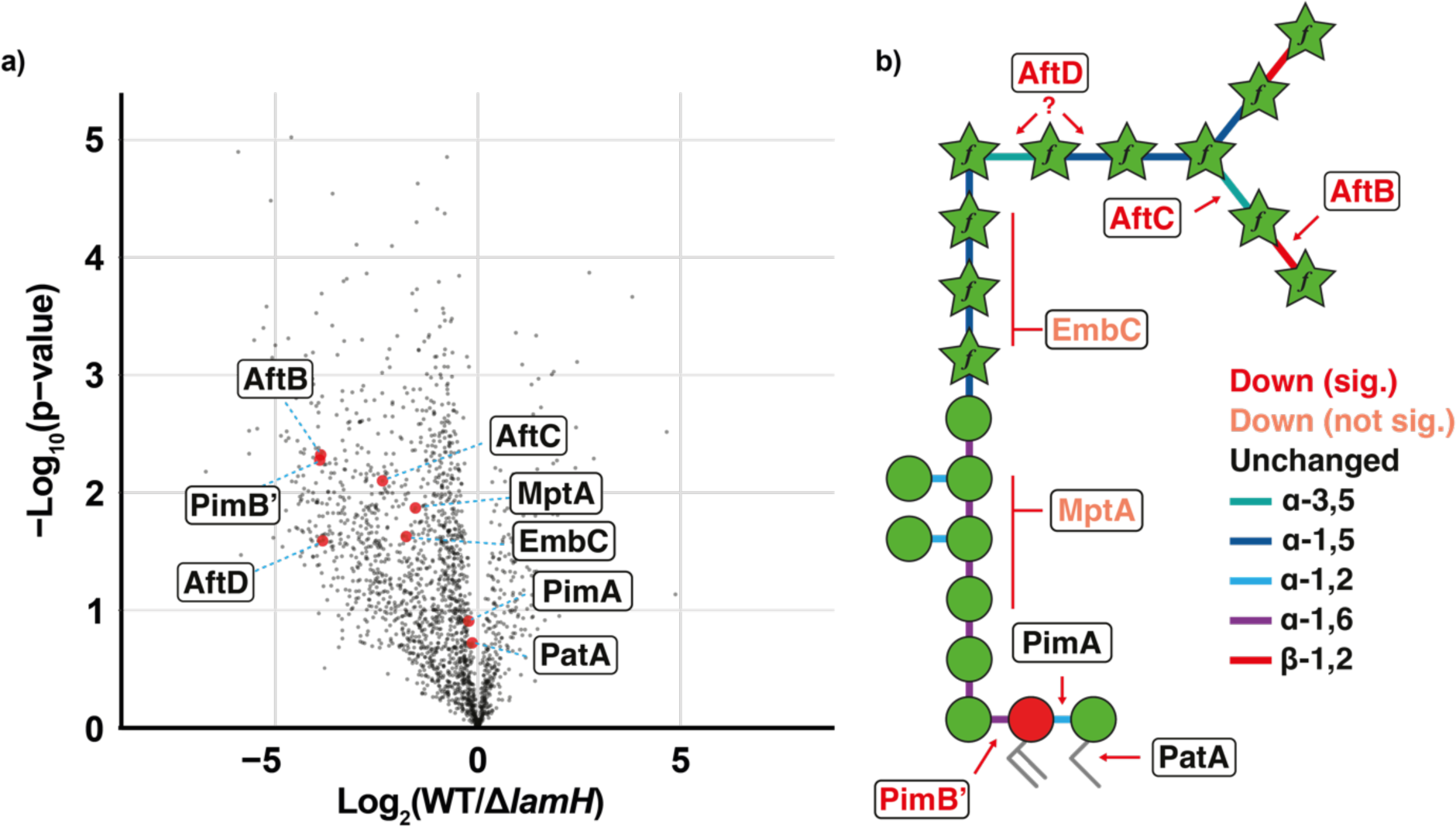
Proteome analysis of *M. bovis* BCG Danish 1331 WT and Δ*lamH*. **a)** Volcano Plot of proteome alterations observed between *M. bovis* BCG and Δ*lamH* visualised as log_2_(Δ*lamH*/WT) and −log_10_(p-values). Proteins of interest in the LAM biosynthetic pathway are denoted in red. Plots showing quantitation of statistically significant proteins are found in Extended Data Fig. 3. The full list of differentially expressed proteins is in Extended Data Table 1. **b)** Eight proteins associated with LAM biogenesis were observed in the proteomics dataset. These have been annotated next to the bond on the simplified LAM structure for which they are understood to generate. Proteins coloured red meet statistical significance for being lower in abundance (Extended Data Fig. 3). Those in orange appear lower in abundance but do not meet statistical significance and those in black have no change associated with their expression.

### LamH is required for efficient transition from lag-phase growth

Given that LAM was recently reported to be involved in septation, we next asked whether loss of LAM turnover would impact growth kinetics^16^. Surprisingly, loss of *lamH* dramatically extended lag-phase growth (Fig. 5a). Under the conditions tested, wild-type bacteria have a lag phase of approximately 5 days. In contrast, the Δ*lamH* mutant extends lag to approximately 14 days (Fig. 5a). This phenotype could be partially complemented with *lamH* but was not complemented by production of the catalytic null variant of the enzyme (Fig. 5a). To understand if *lamH* was involved in both exit from lag-phase and exponential growth, we performed a linear regression on the linear portion of the growth curve (Fig. 5b). This analysis revealed that exponential growth rate of the mutant was reduced, by approximately 20% (Fig. 5b). In contrast to the lag defect and the capsular AM production, this aspect is fully complemented. This suggests that the time point between lag and exponential growth phases is more susceptible to LAM turnover deficit. We interpret these data to suggest that AM acts in part as a molecular cue for division in these bacteria.

**Figure 5.**
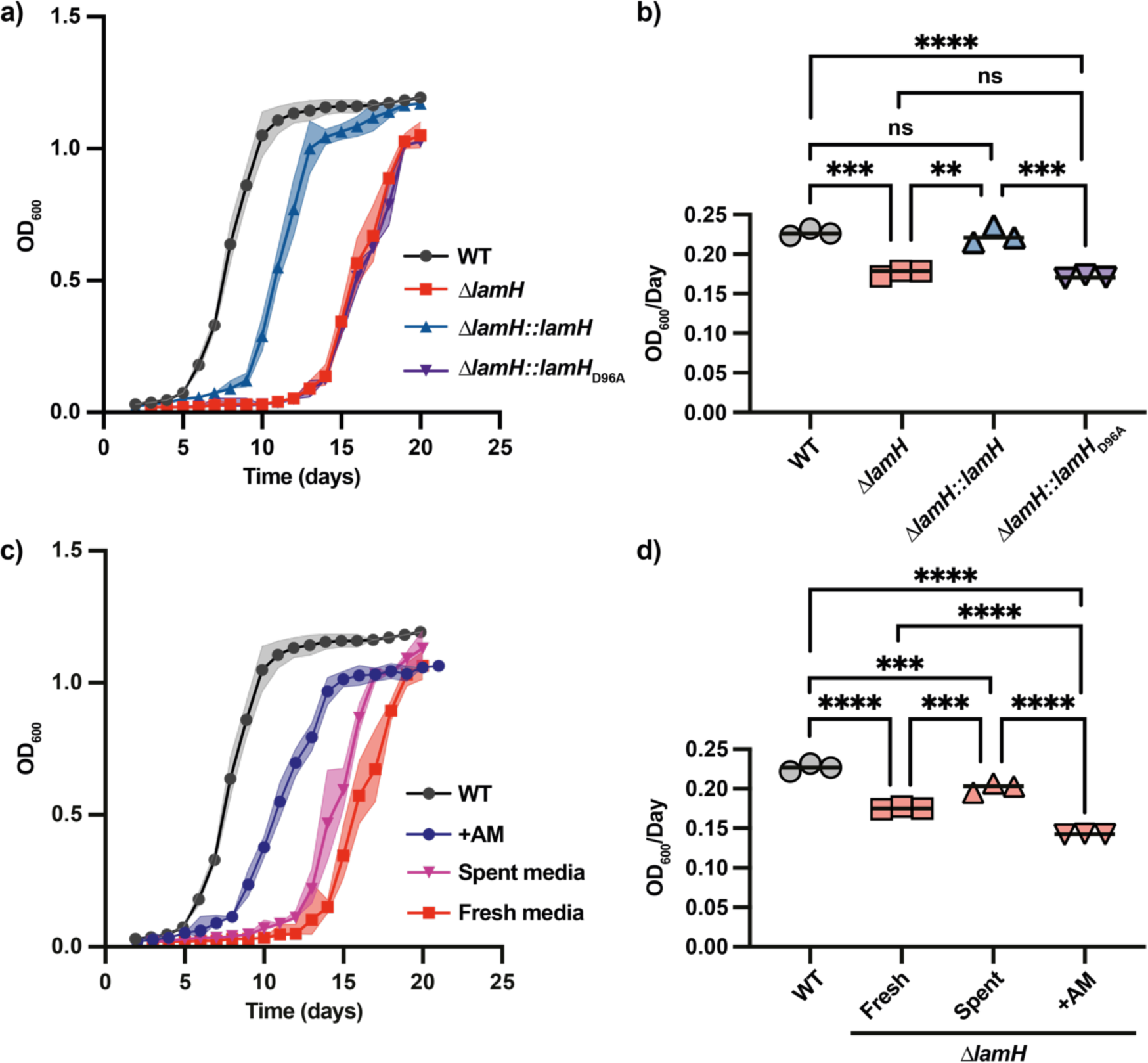
LamH-catalysed AM turnover is required for efficient exit from lag-phase. **a)** Growth kinetics of the indicated strains. n = 3 biological replicates. The shaded area represents the 95% confidence intervals of three biological replicates. **b)** The linear portion of the exponential phase of growth from Fig. 5a was fit to a linear regression and the slope of that line is presented. **c)** The Δ*lamH* strain was grown either in spent 7H9 media from wild-type bacteria, fresh media or media supplemented with 0.5 mg/mL LAM. The shaded area represents the 95% confidence intervals of three biological replicates. The values for WT are included from Fig. 5a for comparison. **d)** The linear portion of the exponential phase of growth from Fig. 5d was fit to a linear regression and the slope of that line is presented. The values for WT are included from Fig. 5b for comparison. For all analysis significance was determined using a one-way ANOVA with Tukey’s post-hoc test. *P < 0.05, **P < 0.01, ***P < 0.001, ****P < 0.0001.

### LamH-derived arabinomannan triggers exit from lag-phase

If capsular AM is a molecular signature for exit from lag phase or growth rate, we would expect that its location in the capsule means that spent media from wild-type bacteria may include sufficient material to alter the growth defects of the Δ*lamH* mutant. We therefore cultivated the mutant in spent media from a mid-exponential wild-type culture. As shown in Fig. 4c, this spent media partially rescued the lag-phase and exponential growth rate defects.

Reasoning that the dose of AM may not be sufficient in spent media, we degraded LAM *in vitro* using recombinant LamH, and added it to fresh media at a concentration of 0.5 mg/mL. This rescued the lag-phase defect to a level similar to genetic complementation (Fig. 4c). The growth rate, however, was significantly lower (Fig. 4d) and the bacteria failed to reach the optical density of the wild type (Fig 4a,c). From this we conclude that *lamH*-driven production of AM acts as a molecular signal for the outgrowth of mycobacteria.

### Reduced expression LamH impairs M. tuberculosis *replication in macrophages*

To assess the impact of *lamH* loss on *M. tuberculosis* host survival we first sought to create a deletion mutant using specialised transduction. Despite repeated attempts we could not generate this mutant. This may be a consequence of the extended lag phase phenotype observed in *M. bovis* BCG or polar effects on nearby genes. To circumvent this, we used CRISPRi to knock-down the expression of *lamH*^42^. Upon addition of anhydrotetracycline (ATc) to the resulting strain Rv0365::pLJR965, the expression of *lamH* was reduced by approximately 80% relative to the same strain grown in the absence of ATc (Extended Data Fig. 5). We also generated a parental strain transformed with an empty plasmid, which upon addition of ATc maintained the expression levels of *lamH* (Extended Data Fig. 5).

We next validated the requirement of *lamH* for intracellular growth within macrophages through the infection of THP-1-derived macrophages with the *lamH* conditional mutant and the parental strain, in the presence or absence or ATc, and determined viable counts 1, 2, and 3 days after infection. Both the *lamH* conditional mutant and the parental strain retained their infective capacity in the absence of ATc (Fig 6a). Conversely, only the *lamH* conditional strain showed reduced viability in the presence of ATc being significantly reduced at all time points after infection (0.38-log reduction at 24 h; 1.01 log-reduction at 48 h; 0.97-log reduction at 72h) (Fig 6b). These results indicate that *lamH* is important for the pathogenesis of mycobacteria.

**Figure 6.**
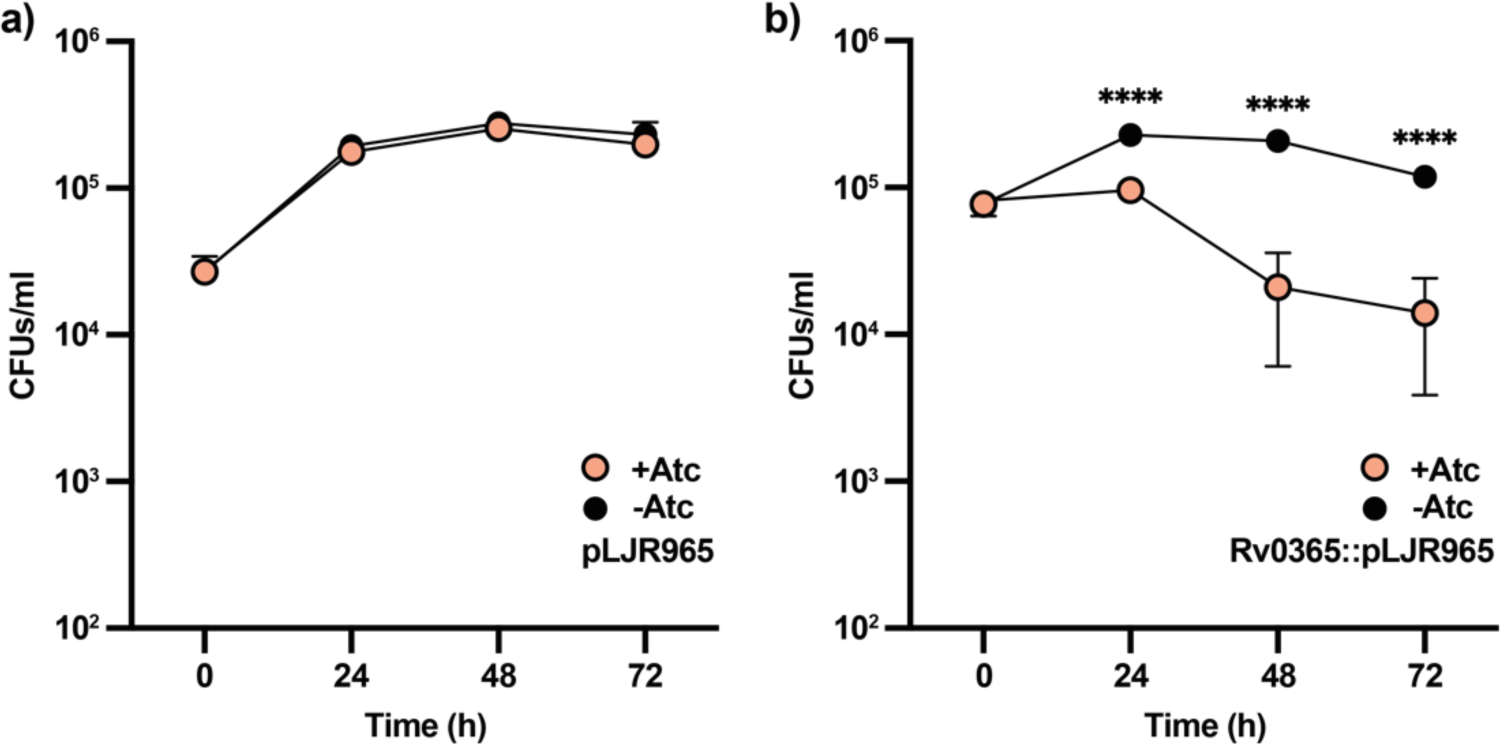
Reduction of *lamH* expression leads to a decrease in fitness in macrophages. **a)** The parental strain including the empty plasmid pLJR965 was used to infect THP-1-derived macrophages in the presence and absence of ATc. Bacterial intracellular viability was examined at the indicated time points. **b)** A strain carrying the *lamH*::pLJR965 plasmid was also used to infect THP-1-derived macrophages in a similar manner. This strain manifested a growth defect in the presence of ATc that was not observed when ATc was omitted from the culture medium. Data are representative of two independent experiments with similar results. Significant differences were calculated by a multiple t-test. ****, *P* < 0.0001.

## Discussion

The activity of a GH76-like α-1,6-mannosidase in mycobacteria was first reported by Yokoyama and Ballou in 1989, and yet more than three decades later the enzyme has eluded characterisation^43^. According to the CAZy database, only 11 GH76 enzymes have been characterised to date, none of which are from pathogenic bacteria^44^. Our biochemical data identify LamH as an LM/LAM hydrolase that is specific for the first α-1,6 linked mannose attached to the AcPIM_2_ anchor of LM/LAM (Fig. 2h). We explored this specificity using structurally homogenous yeast mannan from mutant strains. Our data demonstrate the enzyme will not digest mannan with α-1,2 backbone decorations, which are present on much of the mannan backbone of LAM distal from the AcPIM_2_ anchor^6^. This specificity is possibly driven by the predicted β-hairpin capping the active site (Fig. 1b). Alignment with the Aman6 structure suggests that binding of α-1,6 mannotetraose should be accommodated within the LamH active site, however the capping β-hairpin and active site would clash with substituents on mannose residues in the −1 and −2 positions. Future work will be required to determine the precise substrate recognition mechanism of LamH.

Our biochemical data indicate that AcPIM_2_ is the dominant, if not sole, anchor of LM and LAM in *M. bovis* BCG. This is consistent with prior studies, although we observe no detectable variation in the acylation status of these molecules^45^. Production of AcPIM_2_ upon digestion with LamH suggests a possible pathway for recycling of this lipid anchor into new rounds of LM/LAM and AM biogenesis. This is reminiscent of how mycobacteria have been shown to recycle components of peptidoglycan and trehalose mycolates^46–48^. Given its high specificity, it is likely that LamH will be a useful tool for probing the structure of the PIM anchor of LM/LAMs isolated from other mycobacterial species and critically, clinical isolates. The Jackson laboratory recently reported the identification of an Aman6 isozyme from *Bacillus circulans* for a similar purpose, which demonstrated activity against LAM^49^. However, this enzyme appeared to hydrolyse LM/LAM at more positions than LamH, complicating the analysis of the reaction products.

Three main hypotheses have been put forward to explain how LM, LAM or their by- products are released to the host. First, LM/LAM may be extracted from the cytoplasmic membrane and trafficked to the outside of the cell through an active secretion system. The lipoprotein LprG has been implicated as a potential LM/LAM carrier protein, and loss of *lprG* leads to reduced surface exposure of LM/LAM^50,51^. However, another report suggests that the physiological role of this protein is to transport triacylglycerols^52,53^. Alternative mechanisms for active LM/LAM secretion have yet to be discovered. The second mechanism is through the production of vesicles by the bacteria, which are reported to contain LM/LAM^54,55^. Finally, the production of capsular AM has long been known^20,56^. Indeed, capsular AM has been shown to drive the production of protective antibodies^57,58^. To our knowledge, this work provides the first mechanistic basis for the generation of capsular AM (Fig. 3c). Our data also provide evidence that generation of this material is important for pathogenesis. While we were unable to make a complete deletion of *lamH* in *M. tuberculosis* H37Rv, CRISPRi-mediated silencing reduced its expression by approximately 80%. Despite incomplete knock-down, this strain had an approximately 10-fold reduction of growth in macrophages. This echoes a prior report where expression of *lamH* in *M. smegmatis* mc^2^155 provided a growth-advantage in macrophages^59^. It is therefore possible that LamH-driven AM production is an anti-virulence target.

LM and LAM have important roles during pathogenesis. Yet, all mycobacteria generate LM and LAM, most species of mycobacteria are considered non-pathogenic. Recent work from the Morita identified to a role for LAM in septation, like that observed for lipoteichoic acids in other bacteria^16,60^. Our analysis of Δ*lamH* mutants in *M. bovis* BCG and *M. smegmatis* demonstrate that this enzyme is critical to maintaining LM/LAM homeostasis in the bacteria and that it drives expression of capsular AM. This suggests that the bacteria actively monitor the amount of LAM or AM in the cell and adjust biosynthesis by regulating the levels of synthetic enzymes through some as-of-yet unknown signalling pathway. This hypothesis is supported by the surprising result that loss of *lamH* leads to a profound lag-phase defect in *M. bovis* BCG. Repair of this defect requires the catalytic activity of LamH or the addition of AM to the culture media suggesting that the AM is a signalling molecule. We can rule out the use of AM as a carbon nutrient given that *M. tuberculosis* H37Rv has previously been shown not to be able to utilise D-arabinose or D-mannose as carbon sources^61^. While our data supports a role for AM in a signalling process, it is possible that the signal is not intact AM but is derived from further processing. For example, this could be driven by the action of recently discovered the endo-D-arabinanases^19^. In this context, the abundance of AM or degraded fragments thereof could directly report on cell growth and division, and therefore report on the metabolic state of the bacteria. Taken together, it is tempting to speculate that AM can act like a quorum-sensing molecule in mycobacteria.

## Methods

### Strains and growth conditions

Unless stated otherwise all chemicals and reagents were purchased from Sigma-Aldrich. Strains used were: *Escherichia coli* T7 Shuffle (New England Biolabs) to express Rv0365c, Rv0365c D96A, and BT3792; *S. cerevisiae* Mnn1, Mnn2, and Mnn5^23^; *M. bovis* BCG Danish 1331 WT and *M. bovis* BCG Dan_Δ0378^30^; and *M. smegmatis* mc^2^155 WT^62^ and *M. smegmatis* mc^2^155_Δ0740. *E. coli* strains, up to 100 mL, were grown in lysogeny broth. *E. coli* cultures used for protein purification were grown in terrific broth at 37 °C with agitation. *S. cerevisiae* cultures were grown in yeast extract-peptone-dextrose media at 37 °C with agitation. *M. bovis* BCG Danish 1331 was grown in 7H9 media at 37 °C with 5% CO_2_. *M. smegmatis* mc^2^155 was grown in Middlebrook 7H9 media at 30 °C. *M. tuberculosis lamH* conditional mutant was generated using CRISPRi technology as previously described (Rock et al 2017). Briefly, two oligonucleotides (5’-TGGCTAACAGCTATTACGACTCCC-3’) complementary to Rv0365 were synthesized, annealed, and cloned into pLJR965 plasmid. The selection of the sgRNAs was based on a theoretical degree of repression of 100%, according to the PAM sequences. The vectors were transformed into *E. coli* and extracted to confirm the presence of the sgRNA by Sanger sequencing using the primer, 5’-TTCCTGTGAAGAGCCATTGATAATG-3’. The resulting construct was electroporated into the parental mycobacterial strain H37Rv and selected on 25 μg/ml kanamycin. The parental carrying the empty vector (without a targeting sgRNA) were used as negative controls.

To monitor the growth of *lamH* conditional mutant and the parental strain harbouring empty pLJR965 plasmid, mid-log liquid cultures were diluted to an optical density at 600 nm (OD_600_) of 0.05 in 7H9-ADC + 0.05% tyloxapol with or without 200 ng/mL ATc and incubated in static at 37 °C. OD_600_ was monitored for 15 consecutive days.

### ORBIT-Mediated Mutagenesis of M. smegmatis mc^2^155

Twenty mL of *M. smegmatis* mc^2^155 [pKM461] cells were grown in 7h9 media containing kanamycin at 37 °C in a shaking incubator. At an of OD600 0.5, 500 ng/mL of ATc was added to the culture. This was incubated for a further 3 h at 37 °C. The culture was washed twice in 20 mL of sterile 10% cold glycerol. Following the second wash, cells were collected via centrifugation and resuspended in 2 mL of 10% cold glycerol. On ice, 380μL of the electrocompetent cells, 200 ng of the pKM464 payload plasmid and 1μg of the targeting oligonucleotide (Supplementary Table 2) were added to 2 mm gap width electroporation cuvette. The cells were then electroporated at 2.5kV before overnight incubation (37 °C) in 7h9 media with 0.05% Tween 80. The next day, 0.5 mL of culture was spread onto 7h10 plates containing hygromycin (50 μg/mL) and incubated for 3-4 days at 37°C for the selection of transformants. Selected colonies were analysed by PCR.

### Quantification of mRNA of lamH and nearby genes by qPCR

Cultures were grown to log phase and then diluted back to an OD_600_ of 0.02 in the presence or absence of 200 ng/ml ATc. Target knockdown was allowed to proceed for 4 days. Next, cells were harvested by centrifugation, resuspended in TRIzol (Thermo Fisher), and disrupted by bead beating (Lysing Matrix B, MP Biomedicals). Total RNA was isolated by RNA miniprep (Zymo Research). cDNA was prepared with random hexamers as per manufacturer instructions (Life Technologies Superscript IV). cDNA levels were quantified by quantitative real-time PCR (qRT-PCR) on an Applied Biosystems light cycler (Applied Biosystems) using a SYBR Green PCR Master Mix (Thermo Fisher Scientific) using specific primers. Signals were normalized to the housekeeping *sigA* transcript and quantified by the ΔΔCt method.

### Radiolabelling of mycobacterial lipids

*M. bovis* BCG Danish 1331 cultures were grown until an OD_600_ of 0.2 was reached. Cultures were radiolabelled with addition of 10 µCi/ml acetic acid, sodium salt [1-^14^C] (specific activity 50-62 mCi /mmol (1850-2294 MBq/mmol; Perkin Elmer) and further incubated until an OD_600_ of 0.8 was reached.

### Protein expression and purification

The expression construct for Rv0365c was purchased as codon-optimised constructs from Twist Biosciences in a pET28a vector including residues 2-376 with an N-terminal thrombin cleavage site and hexa-histidine tag. The BT3792 expression plasmid was previously reported^23^. Recombinant proteins were expressed in competent *E. coli* T7 Shuffle cells. Site-directed mutants were generated using the New England Biolabs Q5 mutagenesis kit. For purification of Rv0365c, cultures were grown until an OD_600_ of 0.6 was reached, at this point 0.1 mM IPTG was added to induce protein expression. Cells were further incubated for 16 hours at 14 °C. Cells were harvested by centrifugation at 6000 x *g* for 20 min at 4 °C and resuspended in 100 mM HEPES pH 7.5, 300 mM NaCl, 5 mM imidazole pH 7.5. Resuspended pellet was stored at −20 °C until use. Pellets were thawed and 5% glycerol and 1% Tween20 was added. Additionally, 1 mg/mL deoxyribonuclease from bovine pancreas (Sigma-Aldrich) was added to the resuspension and incubated on ice for 30 min. The cell resuspension was lysed with three passages through a French pressure cell. Cell debris was pelleted by centrifugation at 40,000 x *g* for 45 min at 4 °C. Enzymes were purified using immobilised metal affinity chromatography (IMAC) on nickel Sepharose resin in a gravity column. Bound protein was eluted from the column with increasing concentrations, 5 mM to 500 mM of imidazole washes. Positive fractions were determined by SDS-PAGE and dialysed into 100 mM HEPES pH 7.5, 300 mM NaCl buffer at 4 °C. The protein was concentrated to a final volume of 500 µL using a 30 kDa molecular weight cut off protein concentrator (Thermo Scientific).

For purification of BT3792, cultures were grown until an OD_600_ of 0.6 was reached. Protein expression was induced with the addition of 0.2 mM IPTG, and cultures were incubated for a further 16 h at 16 °C with 180 rpm shaking. As before, cell pellets were harvested by centrifugation at 6000 x *g* for 20 minutes at 4 °C and resuspended in 150 mM Tris pH 8.0, 300 mM NaCl, 20 mM imidazole pH 8.0. Cells were stored at −20 °C until use. The pellets were thawed and 1 mg/mL deoxyribonuclease from bovine pancreas (Sigma-Aldrich) was added to the resuspension and incubated on ice for 30 min. Cells were lysed with three passages through a French pressure cell, and cell debris was pelleted by centrifugation at 40,000 x *g* for 45 min at 4 °C. Enzymes were purified by IMAC using nickel Sepharose resin in a gravity column. Bound enzymes were eluted from the column with an imidazole gradient ranging from 20 mM to 500 mM. Positive fractions were identified by SDS-PAGE and dialysed for 16 h at 16 °C into 150 mM Tris pH 8.0, 300 mM NaCl buffer. As before, protein was concentrated to a final volume of 500 µL using a 30 kDa molecular weight cut off protein concentrator (Thermo Scientific).

### Enzyme assays

Purified protein, at a final concentration of 1 µM, was incubated with 1 mg / ml substrate at 37 °C for 16 h (unless stated otherwise) in 100 mM HEPES buffer pH 7.5. Reactions were heat inactivated by incubation at 100 °C for 10 mins and analysed by thin-layer chromatography.

### Thin-layer chromatography (TLC)

Samples were spotted onto a TLC plate (Merck, TLC Silica Gel 60 F_254_), and separated until the solvent front had reached 5 mm from the top of the sheet. The TLC plate was dried, and either stained and heated or exposed to X-ray film for radioactive samples.

### TLC solvent systems

For the separation of mannans, samples were separated in n-butanol: acetic acid: water (2:1:1 v/v/v), stained with orcinol and charred to reveal products. For the analysis of LAM, samples were separated in chloroform: methanol: water (65:25:3 v/v/v), stained with either orcinol or MPA, and charred by heating. Analysis of PIMs was performed by two-dimensional TLC. The samples were separated in the first direction in chloroform: methanol: water (60:30:6 v/v/v) and in the second direction in chloroform: acetic acid: methanol: water (65:25:3:6 v/v/v/v). TLCs were annotated as per Driessen *et al.,* (2009)^63^.

### Purification of β-*mannan from* S. cerevisiae

Purification of β-mannan from *S. cerevisiae* was performed as previously described^23^. Eight litres of the desired *S. cerevisiae* strain were grown at 37 °C for 24 h. Pellets were harvested by centrifugation at 6000 x *g* for 20 min. The pellets were then pooled and stored at −20 °C. The pooled cell pellets were resuspended in 20 mL 0.02 M citrate buffer pH 7.0 and then autoclaved for 90 min at 121 °C. The sample was centrifuged at 6000 x *g* for 10 min, and the supernatant was collected. The remaining pellet was resuspended in 75 mL of 0.02 M citrate buffer pH 7.0 and autoclaved once more at 121 °C for 90 min. The sample was centrifuged at 6000 x *g* for 10 min, and the resulting supernatant was pooled with that previously collected. A volume of 2x Fehling’s reagent, equal to that of the supernatant, was measured out and heated to 40 °C. The supernatant was carefully added to the 2x Fehling’s reagent and stirred vigorously at 40 °C for 1 h. Following this, the mixture was centrifuged at 6000 x *g* for 10 min to harvest the pellet. The pellet was dissolved in 8 mL of 3 M hydrochloric acid before 100 mL of methanol: acetic acid solution (8:1 v/v) was added and stirred for 1 h at room temperature. The mixture was centrifuged at 12,000 x *g* for 15 min and the pellet was collected. The pellet was resuspended in 20 mL methanol: acetic acid solution (8:1 v/v) and centrifuged again. This process was repeated until the pellet was colourless. The pellet was the left to dry at room temperature overnight. The dry pellet was resuspended in 20 mL dH_2_O and then dialysed against 4 L of dH_2_O for 24 hours. The mixture was lyophilised to complete dryness.

### Purification of mycobacterial glycolipids

One litre of mycobacterial culture was grown until an OD_600_ of 0.8 was reached and cells were harvested by centrifugation at 6000 x *g* for 10 min. The pellet was resuspended in 20 mL PBS, 0.1% Tween-80, and lysed by bead-beating. The cell lysate was transferred to a Teflon-capped glass tube and an equal volume of phenol was added. The mixture was heated to 85 °C and incubated for 2 h with regular mixing. The aqueous phase was separated from the phenol phase by centrifugation at 4000 x *g* for 10 min and transferred to a fresh tube. The phenol wash is repeated twice more. The glycan mixture was dialysed exhaustively against tap water overnight. Following this, the mixture was further dialysed against ddH_2_O for 2 h and then lyophilised.

### Polar lipid extraction

Mycobacterial cultures were grown until an OD_600_ of 0.8 was reached and harvested by centrifugation at 6000 x *g* for 10 min. The pellets were resuspended in PBS, 0.1% Tween-80, and transferred to a Teflon-capped glass tube. 2 mL methanol: 0.3 % NaCl (100:10 v/v) and 2 mL petroleum ether (60-80) was added to the pellet and mixed for 24 h on a rotator. The sample was centrifuged at 4000 rpm for 10 min to form a bilayer, and the upper layer was transferred to a fresh tube. An additional 2 mL of petroleum ether is added to the remaining lower layer and further mixed for 1 hour. The sample was centrifuged as before, and the upper layers pooled. The petroleum ether extracts were dried under a stream of nitrogen to give the non-polar lipids. To the remaining lower layer, 750 µL of chloroform: methanol: 0.3% NaCl (9:10:3 v/v/v) was added and mixed for 2 h. The sample was centrifuged at 4,300 rpm for 10 min and the supernatant was transferred to a fresh tube. The pellet was resuspended in 950 µL chloroform:methanol:0.3% NaCl (5:10:4 v/v/v) and mixed for 30 min. The sample was centrifuged at 3,500 rpm for 5 min and the supernatant was pooled with that from the previous step. The polar lipids were extracted by addition of 1 mL chloroform and 1 mL 0.3 % NaCl to form a bilayer. The lower, polar lipid containing, layer was transferred to a fresh tube and dried under a stream of nitrogen.

### Capsular polysaccharide extraction and analysis

*M. bovis* BCG Danish cultures were grown on 7H10 agar plates for 4 weeks. Subsequently, cells were scraped from the plates and a cell mass was measured. The cells were then resuspended in ddH_2_O and gently vortexed for 1 minute to shed capsular material. Cells were harvested by centrifugation at 2000 x *g* for 10 minutes. The capsule-containing supernatant was collected and filtered using a 0.45 µM Millipore filter to ensure no bacterial debris remained. The capsular extract was frozen and lyophilised to complete dryness.

Fluorescent labelling of the capsular polysaccharide extracts was carried out following the methods of Ruhaak *et al.,* (2010)^64^. In brief, 50 µl of capsular material was mixed with 25 µl of freshly prepared label (48 mg/mL 2-aminobenzamide (Ludger) in DMSO/acetic acid (85:15 v/v)). Next, 25 µl of 1 M 1-picolane-borane in DMSO was added to achieve a final volume of 100 µl. Subsequently, the mixture was incubated at 65 °C for 2 hours. The samples were then allowed to cool to room temperature before being analysed.

Analysis of the fluorescently labelled capsular extracts was performed using a Phenomenex BioZen 1.8 µM size exclusion chromatography-2 column (300 x 4.6 mm, 00H-4769-E0) at room temperature on a Dionex Ultimate 3000 uHPLC. Mobile phase was 0.1 M phosphate buffer pH 6.8 with 0.025 % sodium azide, flow rate = 0.400 mL / min. Glycans detected with fluorescence detection (Ex = 320 nm Em = 420 nm), sample volume injected = 2.5 µL. Peak identities were confirmed by digestion with α-amylase (Sigma-Aldrich), BT3792, or GH183*_Mab_*.^19,23^

### Growth point determination

Bacterial growth was recorded by taking OD_600_ readings daily, using a portable spectrophotometer until stationary phase was reached. Cultures were grown static, in triplicate, in 7H9 media at 37 °C with 5 % CO_2_ using Corning culture flasks (Sigma-Aldrich, CLS431082). Starter cultures were grown until mid-log phase (OD_600_ = 0.6) was reached and subsequently diluted to an OD_600_ of 0.01 in 10 mL 7H9 media, and growth was recorded every 24 h.

### Sample preparation for proteomic analysis

Acetone precipitated proteome samples were solubilized in 4% SDS, 100 mM Tris pH 8.5 by boiling for 10 min at 95 °C. The protein concentrations were assessed using a bicinchoninic acid protein assay (Thermo Fisher Scientific) and 100 μg of each biological replicate prepared for digestion using Micro S-traps (Protifi, USA) according to the manufacturer’s instructions. Samples were reduced with 10mM DTT for 10 mins at 95°C and then alkylated with 40 mM IAA in the dark for 1 h. Samples were acidified to 1.2% phosphoric acid, diluted with seven volumes of S-trap wash buffer (90% methanol, 100 mM tetraethylammonium bromide pH 7.1), then loaded onto S-traps and washed 3 times with S-trap wash buffer. Samples were then digested overnight with Trypsin (1:100 protease:protein ratio, Solu-Trypsin, Sigma) within 100 mM Tetraethylammonium bromide pH 8.5. Following overnight digestion, peptides were collected by centrifugation using washes of 100mM Tetraethylammonium bromide, followed by 0.2% formic acid followed by 0.2% formic acid / 50% acetonitrile. Samples were dried down and further cleaned up using C18 Stage tips to ensure the removal of any particulate matter^65,66^.

### Reverse phase liquid chromatography–mass spectrometry

Proteome samples were re-suspended in Buffer A* (0.1% trifluoracetic acid, 2% acetonitrile) and separated using an Ultimate 3000 UPLC (Thermo Fisher Scientific) equipped with a two-column chromatography set up composed of a PepMap100 C18 20 mm x 75 μm trap and PepMap C18 500 mm x 75 μm analytical column (Thermo Fisher Scientific). Samples were concentrated onto the trap column at 5 μL/min for 6 minutes with Buffer A (0.1% formic acid, 2% DMSO) and then infused into a Orbitrap Fusion™ Eclipse™ Tribrid™ mass spectrometer equipped with a FAIMS Pro interface (Thermo Fisher Scientific) at 300 nl/minute via an analytical column. 89-minute analytical runs were undertaken by altering the buffer composition from 2% Buffer B (0.1% formic acid, 77.9% acetonitrile, 2% DMSO) to 28% B over 70 minutes, then from 28% B to 40% B over 9 minutes, then from 40% B to 80% B over 3 minutes. The composition was held at 80% B for 2 minutes, and then dropped to 2% B over 0.1 minutes before being held at 2% B for another 2.9 minutes. Each biological replicate was analysed using four different FAIMS compensation voltages (CVs, −25, −35, −45 and −65) in a data-dependent manner switching between the acquisition of a single Orbitrap MS scan (450-2000 m/z, maximal injection time of 50 ms, an AGC set to a maximum of 1*10^6^ ions and a resolution of 60k) every 3 seconds followed by Orbitrap MS/MS HCD scans (using the “Auto” mass range setting, a NCE of 25;32;40%, maximal injection time of 120 ms, an AGC set to a maximum of 500% and a resolution of 30k) as well as a Orbitrap EThcD scan (NCE 25%, maximal injection time of 120 ms with an AGC of 500% and a resolution of 30,000) undertaken on each precursor.

FragPipe version 19^67–71^ was used to process the resulting proteome dataset with proteins identified by searching against the *M. bovis* BCG Danish 1331 proteome (NCBI accession: CP039850.1) allowing carbamidomethyl (57.0214Da) of cystines as a fixed modification as well as oxidation of methionine (15.9949Da), N-terminal acetylation (42.0106Da) in addition to hexose (162.0528Da) and 2*Hexose (324.1056Da) on serine and threonine residues as variable modifications. The resulting datasets were filtered using the default FragPipe parameters of 1% peptide/protein level false discovery rates. IonQuant was utilised for quantitative proteome comparisons with matching between runs enabled across biological replicates using the default parameters. The resulting combined MSfragger protein level output was process using Perseus (version 1.6.0.7) with missing values imputed based on the total observed protein intensities with a range of 0.3 σ and a downshift of 1.8 σ^72^. Statistical analysis was undertaken in Perseus using two-tailed unpaired t-tests.

## Data availability

The mass spectrometry proteomics data has been deposited in the Proteome Xchange Consortium via the PRIDE partner repository with the data set identifier: PXD042653 (Username: Password: dTg3p1d7)^73^.

## Acknowledgements

This work was supported by the Biotechnology and Biosciences Research council (grant BB/S010122/1 and BB/X00841X/1 to PJM, and BB/X016749/1 to ECL). We are also grateful to the Wellcome Trust for supporting this work (226644/Z/22/Z to PJM, S.W., and E.L., 209437/Z/17/Z to A.L.L., and a studentship to S.B.) G.S.B is supported by the Medical Research Council (MR/S000542/1). This work was supported in part by the MICINN grant PID2022-138694OB-I00 and the NIH grant R01GM148075 to M.E.G. We thank the Australian Research Council for a Future Fellowship to NES (FT200100270) and ARC Discovery Project Grants to NES and SJW (DP210100362, DP210100233, DP210100235). We thank the Melbourne Mass Spectrometry and Proteomics Facility of The Bio21 Molecular Science and Biotechnology Institute for access to MS instrumentation. We thank members of the Birmingham mycobacteriology group for helpful discussions.

---

**Supplementary Table 1. DDA protein level LFQ analysis of *M. bovis* BCG Danish Δ*lamH* compared to WT.** The Perseus processed MSfragger search results for the protein analysis of three biological replicates of strains WT and Δ*lamH* are provided. For each identified protein, the log2 LFQ protein values, the t-test information including the −log_10_(*p*-value), difference in the mean between the groups and if the resulting *p*-values is below the multiple hypothesis corrected *p*-value are provided. For each protein the protein score, number of peptides identified, and protein length are provided.

