## Extended Data for "The mycobacterial glycoside hydrolase LamH enables capsular arabinomannan release and stimulates growth"

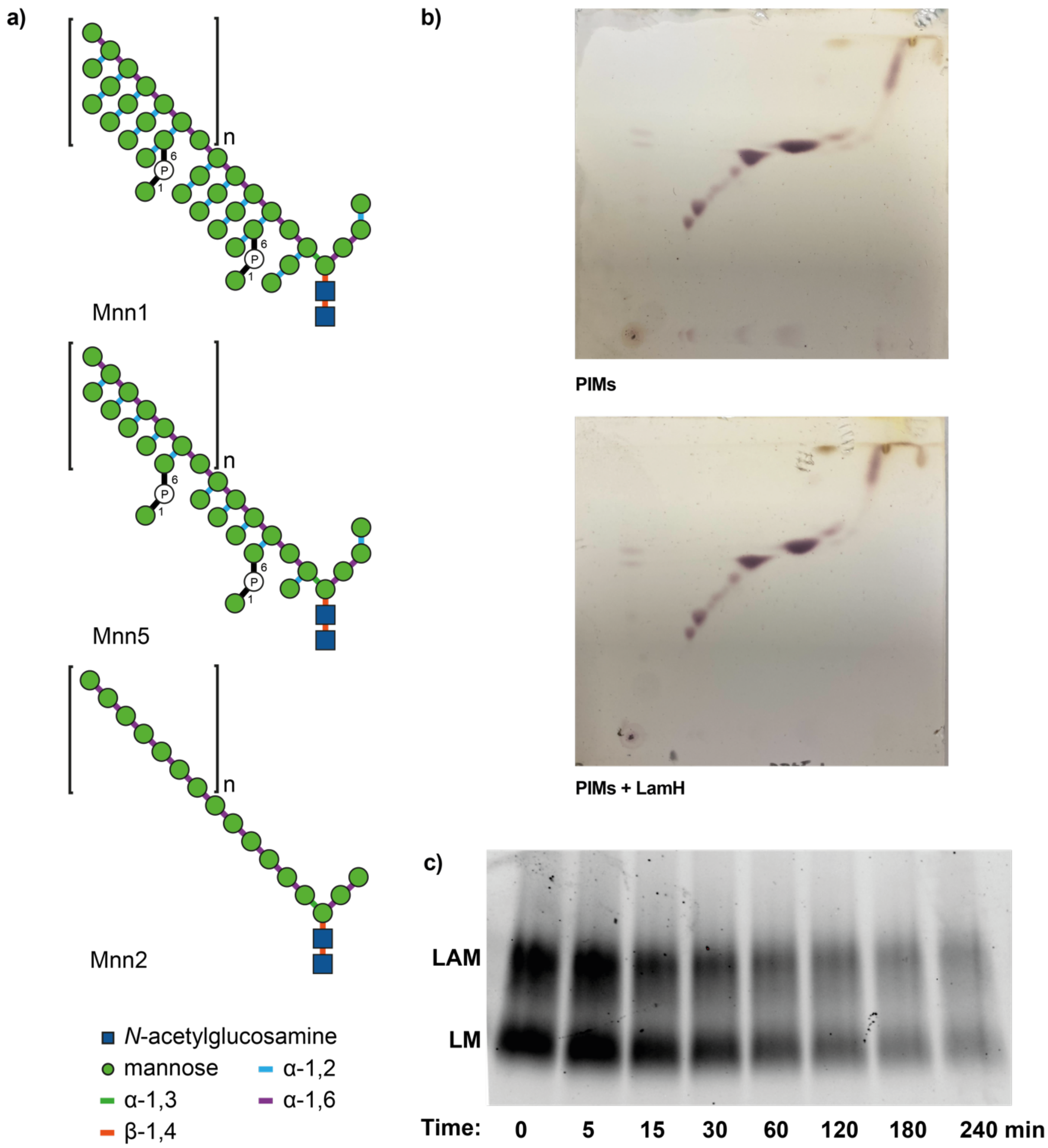


**Extended Data Figure 1. LamH substrate specificity. A)** Structure of the mannans purified from mutant strains of *S. cerevisiae*. **B)** Representative two-dimensional TLC analysis of polar lipids from *M. bovis* BCG Danish wild-type with and without addition of LamH. Equal volumes were loaded onto the TLC and separated in the first direction in solvent system E1: chloroform:methanol:water (60:30:6 v/v/v) and in the second direction in solvent system E2: chloroform:acetic acid:methanol: water (40:25:3:6 v/v/v/v). TLCs were visualised by staining with oricinol and charring. **C)** Example SDS-PAGE time-course data for Fig. 2d. Purified LAM/LM was incubated with LamH at 37 °C and aliquots were taken at the given time points. The reaction mixture was heat inactivated at 100 °C for 10 min. Subsequently, the time points were separated by SDS-PAGE and then stained with Pro-Q Emerald. The gel was visualised by fluorescence imaging at 300 nm on a Bio-Rad Gel Doc XR+. Bands were quantitated using the Bio-Rad ImageLab v6.1 software package and these data are presented in Fig. 2d.

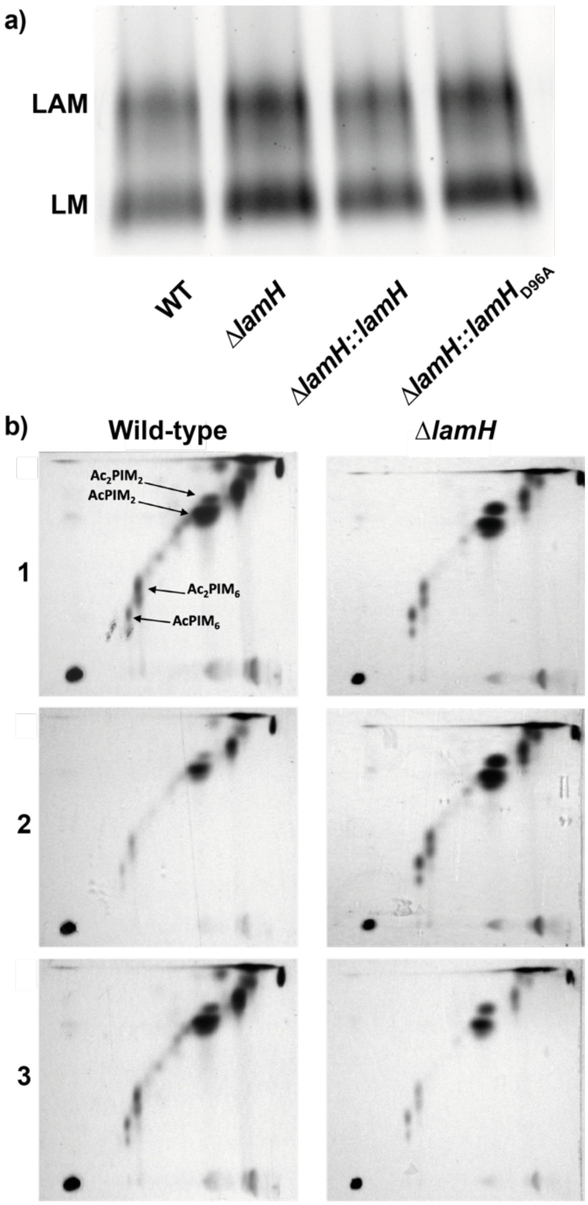


**Extended Data Figure 2. Rv0365c substrate specificity. a)** Representative LM/LAM gel from selected strains of *M. bovis* BCG Danish. The gel was imaged as described above. **b)** Two-dimensional TLC analysis of $^{14}C$ labelled polar lipids from *M. bovis* BCG Danish wild-type and Δ*lamH* mutants. Equal counts of extract were loaded onto the TLC and separated in the first direction in solvent system E1: chloroform:methanol:water (60:30:6 v/v/v) and in the second direction in solvent system E2: chloroform:acetic acid:methanol: water (40:25:3:6 v/v/v/v). TLCs were visualised by exposure to X-ray film by autoradiography. PIMs are annotated as per Driessen *et al.,* (2009). Three biological replicates are presented.

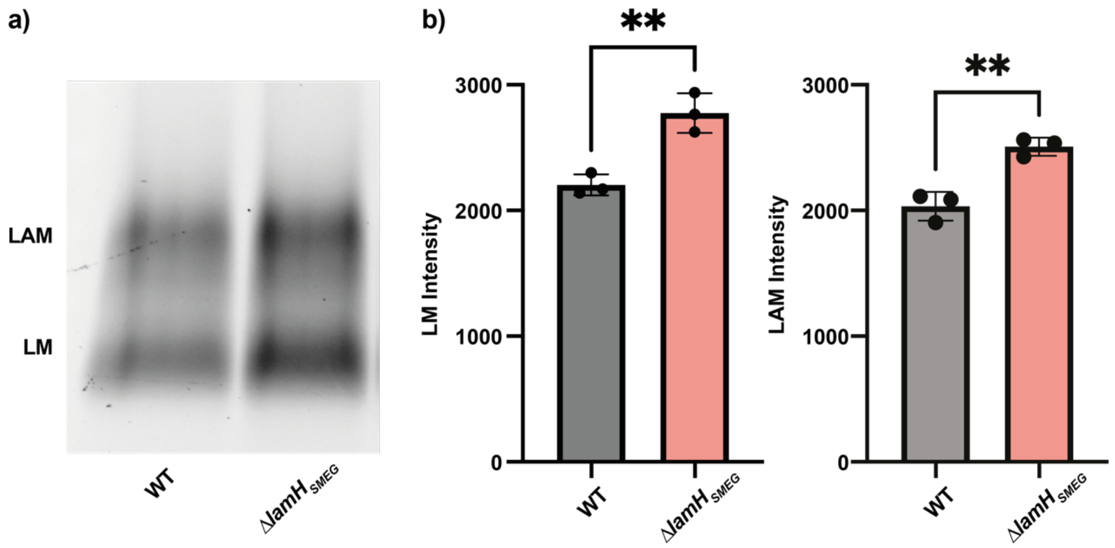


**Extended Data Figure 3. LamH function is conserved in *M. smegmatis*. a)** LAM/LM purified from an equal number of cells was analysed by SDS-PAGE and Pro-Q Emerald staining. The gel was visualised by fluorescence at 300 nm on a Bio-Rad Gel Doc XR+. **b)** Fluorescence was determined for three biological replicates for both LM and LAM using Bio-Rad ImageLab v6.1 software package and is presented with error bars indicating standard deviation. Significance determined by an unpaired t-test. **$P < 0.01$.

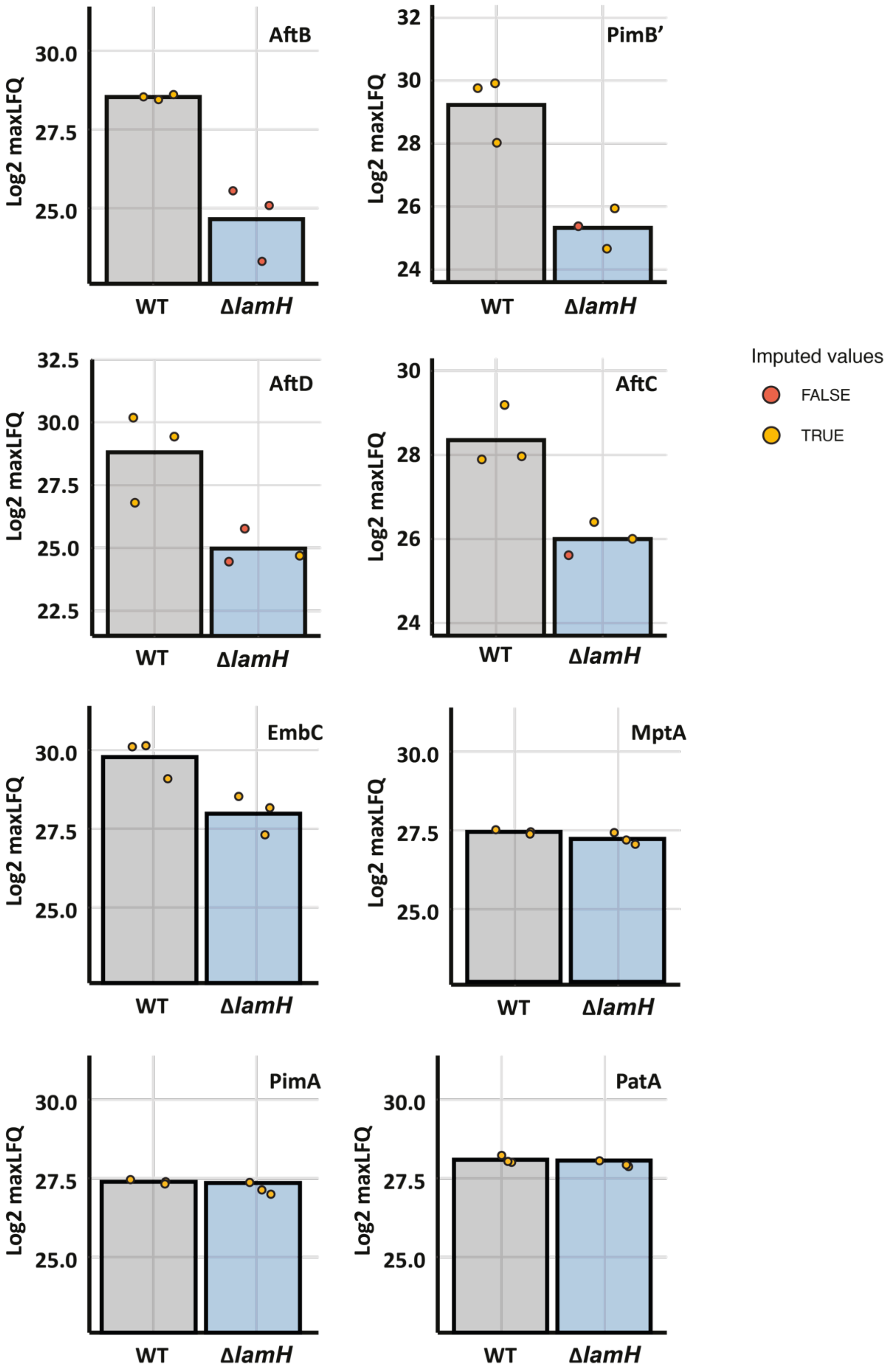


**Extended Data Figure 4. Quantitation of Δ*lamH* proteomics.** Quantitation of individual protein levels observed across biological replicates support the reduction in abundance of proteins associated with LAM biosynthetic pathway within Δ*lamH*. Imputed values are denoted in red while experimental observed values are denoted in orange.

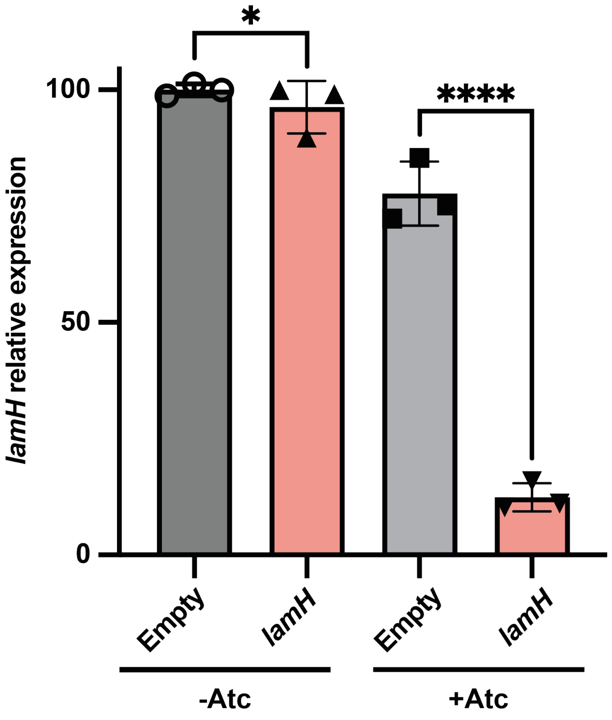


**Extended Data Figure 5. CRISPRi-induced knock-down of *lamH* expression in *M. tuberculosis* H37Rv.** Quantification (mean ± S.E.M., n = 3 biological replicates) of *lamH* mRNA levels by RT-qPCR. Strains were grown ±ATc for ~3 generations before collecting RNA. Significance determined by an unpaired t-test; *$P < 0.05$, ****$P < 0.0001$.
